# Surfactants or scaffolds? RNAs of different lengths exhibit heterogeneous distributions and play diverse roles in RNA-protein condensates

**DOI:** 10.1101/2022.11.09.515827

**Authors:** Ignacio Sanchez-Burgos, Lara Herriott, Rosana Collepardo-Guevara, Jorge R. Espinosa

## Abstract

Biomolecular condensates, thought to form via liquid–liquid phase separation of intracellular mixtures, are multicomponent systems that can include diverse types of proteins and RNAs. RNA is a critical modulator of RNA-protein condensate stability, as it induces an RNA-concentration dependent reentrant phase transition—increasing stability at low RNA concentrations and decreasing it at high concentrations. Beyond concentration, RNAs inside condensates can be heterogeneous in length, sequence, and structure. Here, we use multiscale simulations to understanding how different RNA parameters interact with one another to modulate the properties of RNA-protein condensates. To do so, we perform residue/nucleotide-resolution coarse-grained Molecular Dynamics simulations of multicomponent RNA-protein condensates containing RNAs of different lengths and concentrations, and either FUS or PR_25_ proteins. Our simulations reveal that RNA length regulates the reentrant phase behaviour of RNA-protein condensates: increasing RNA length sensitively rises the maximum value that the critical temperature of the mixture reaches, and the maximum concentration of RNA that the condensate can incorporate before beginning to become unstable. Strikingly, RNA of different lengths are organised heterogeneously inside condensates, which allows them to enhance condensate stability via two distinct mechanisms: shorter RNA chains accumulate at the condensate’s surface acting as natural biomolecular surfactants, whilst longer RNA chains concentrate inside the core to saturate their bonds and enhance the density of molecular connections in the condensate. Using a patchy particle model, we demonstrate that the combined impact of RNA length and concentration on condensate properties is dictated by the valency, binding affinity, and polymer length of the various biomolecules involved. Our results postulate that diversity on RNA parameters within condensates allows RNAs to increase condensate stability by fulfilling two different criteria: maximizing enthalpic gain and minimizing interfacial free energy; hence, RNA diversity should be considered when assessing the impact of RNA on biomolecular condensates regulation.

## I. INTRODUCTION

Intracellular organisation represents a fundamental aspect of regulation with respect to both structure and function. Whilst membrane-bound organelles are responsible for forming large, often permanent compartments within the cell, a more dynamic compartmentalisation can be also achieved through membraneless organelles [1]. Membraneless organelles, also referred to as biomolecular condensates, possess the two key properties of intracellular compartments: the existence of a defined boundary between the compartment and its surroundings, and the ability of components to diffuse freely within the compartment [2, 3]. Biomolecular condensates are thought to form via liquid-liquid phase separation (LLPS) of intracellular mixtures (e.g. proteins, RNA, DNA, and chromatin). Thus, the boundary of condensates is not a traditional lipid membrane, but rather the liquid–liquid interface separating a condensed liquid from its surrounding cytoplasm or nucleoplasm. Since the discovery of P-granules condensates in 2009 [4], important examples of biomolecular condensates including the nucleolus [5, 6], Cajal bodies [7, 8], paraspeckles [9, 10], stress granules [11, 12] and chromatin [13, 14] (a finding which drove the paradigm shift away from the once prominent theory of the 30nm fibre [15–17]) have been exhaustively investigated.

RNA-binding proteins are common components of intracellular biomolecular condensates [18–22]. Several features of RNA-binding proteins underpin their ability to form condensates that are sensitively regulated by RNA. For instance, RNA-binding proteins are multidomain multivalent molecules, many of which can establish sufficiently strong homotypic interactions to act as scaffolds in biomolecular condensates—–e.g., the heterogeneous nuclear ribonucleoprotein 1 (hn-RNPA1) [12, 23], fused in sarcoma (FUS) [24–26], the GTPase-activating protein SH3 domain–binding protein 1 (G3BP1) [27–30] and the Trans-activation response DNA-binding protein 43 (TDP-43) [31–33]. In addition, RNA-binding proteins can bind to RNA both specifically and promiscously via their RNA-recognition motifs (RRMs), positively charged domains (PCDs), and intrinsically disordered regions (IDRs) with low-complexity amino acid sequences [34–36]. Both in vitro and, more recently, in silico experiments have demonstrated the role of RNA as a critical regulator of RNA-protein condensates [37–44]. The RNA-binding protein Whi3 has been shown to partition into different condensates depending on the secondary structure of the RNA to which it is bound [45]. Proteins such as FUS remain soluble in the nucleus, where RNA concentration is high, but form aggregates in the cytoplasm where RNA concentration is lower [46]. Such impact of RNA concentration on protein aggregation may be relevant to rationalize the presence of pathological FUS aggregates (characteristic of Amyotrophic Lateral Sclerosis (ALS)) in the cytoplasm of post-mortem tissues [47] *versus* their absence from the nuclei [48]. The pattern of low levels of RNA promoting condensation *versus* higher concentrations promoting dissolution is described as RNA-driven reentrant phase behaviour [37, 43, 46] and is particularly important when considering the role of RNA in complex coacervation [44, 49–51]. Complex coacervation frequently enables the phase separation of so-called ‘cognate’ proteins which, unlike FUS, cannot sustain LLPS through protein-protein interactions alone, but instead rely on interactions with a partner biomolecule such as RNA [52]. The 25-repeat proline-arginine peptide (PR_25_) is a representative example of a protein which undergoes complex coacervation at physiological conditions driven mostly by electrostatic interactions with RNA [44, 51].

RNA length has been shown to mediate condensate reentrant phase behaviour [53]. Specifically, transcriptional condensate formation and dissolution was shown to be regulated by both RNA length and concentration: short, nascent RNAs present at transcription initiation stimulate condensation, whilst longer nucleic acids resulting from transcriptional bursts promote dissolution [53]. The nature of transcriptional bursting, with the total number of RNA molecules as well as their lengths increasing [54], means it is unclear whether the reentrant phase behaviour of transcriptional condensates is a function of RNA length, concentration, or a combination of both. In vitro experimentation has proved valuable in demonstrating the various ways in which RNA regulates LLPS of RNA-binding proteins [18–22, 37–41, 46]. Complementary, computational modelling and simulations can provide mechanistic insight into the experimental observations, and molecular detail regarding the condensate’s thermodynamic, kinetic and structural properties [55]. Computer simulations can also elucidate condensate properties such as droplet surface tension, protein molecular contact maps, or protein/RNA/DNA conformational ensembles [56–61]. Moreover, key features of LLPS, such as valency [62, 63], topology [64, 65] or binding affinity [66–68] can be precisely controlled. In that sense, Molecular Dynamics (MD) simulations have proved useful for the study of biomolecular condensates at various levels of resolution: ranging from atomistic force fields to lattice-based physical models [16, 51, 69–72].

Here, we use MD simulations, taking advantage of the benefits of computational modelling [56–61], to investigate in molecular detail the role of RNA length and concentration in the regulation of biomolecular condensates. Specifically, we aim to determine how RNA concentration and RNA length cooperate or compete to affect the RNA-dependent reentrant phase behaviour of RNA-binding proteins [18–20, 37, 38, 44, 46]. We use our sequence-dependent Mpipi model [73], which predicts protein phase diagrams in quantitative agreement with experiments, to study the reentrant phase behaviour of two archetypal proteins, FUS and PR_25_, known to undergo LLPS either by homotypic interactions or complex coacervation respectively [74]. Further, we investigate the effect of RNA length on condensate organisation by simulating proteins with mixtures of RNA of different lengths. In particular, we aim to identify whether patterns similar to those previously described with colloidal scaffold-surfactant models [63] can also exist in RNA-protein systems. Finally, by means of a minimal coarse-grained (CG) model [64], we prove that the observed RNA-length-and-concentration-dependent phase behaviour of RNA-protein condensates can also be observed in colloidal particles, and this is strongly regulated by molecular valency, binding affinity and polymer length.

## II. MODELS AND SIMULATION METHODS

Since the formation of phase-separated condensates entails the collective interactions among thousands of different proteins and other biomolecules, the study of LLPS has benefited from the development and application of coarse-grained approaches, including mean field simulations [75–79], lattice-based models [80– 83], minimal models [65, 66], and residue-resolution simulations [57, 58, 67, 84–88]. In this article, we employ two protein/RNA coarse-grained models of distinct resolutions previously developed by us: (1) the residue/nucleotide-resolution Mpipi force field for proteins and RNA [73]; and (2) the MD-Patchy model in which whole proteins are represented as patchy particles, and RNA as self-avoiding flexible polymers [44, 64, 89].

Within the Mpipi force field [73], amino acids and RNA bases are represented by single beads with unique chemical identities (Fig 1(a)) in which hydrophobic, *π*–*π*, and cation–*π* interactions are modelled through a Wang-Frenkel (mid-range) potential [90], and electrostatic interactions via Yukawa/Debye-Hückel (long-range) potentials [84]. Bonded interactions between consecutive residues within the same protein (or nucleotides within the same RNA strand) are described with a harmonic potential. Furthermore, within this model, intrinsically disordered regions of proteins and RNA strands are treated as fully flexible polymers, whilst globular domains are described as rigid bodies based on their corresponding crystal structures taken from the Protein Data Bank (PDB) and adapted to the model resolution. The interactions between ‘buried’ amino acids within globular domains are scaled down by 70% as proposed in Refs. [73, 74]. Finally, the physiological concentration of monovalent ions in solution (i.e., ∼ 150 mM NaCl) is approximated by the screening length of the Yukawa/Debye-Hückel implicit-solvent model [84]. Further details on the force field parameters and simulation setups are provided in the Supplementary Material (SM).

**FIG. 1:**
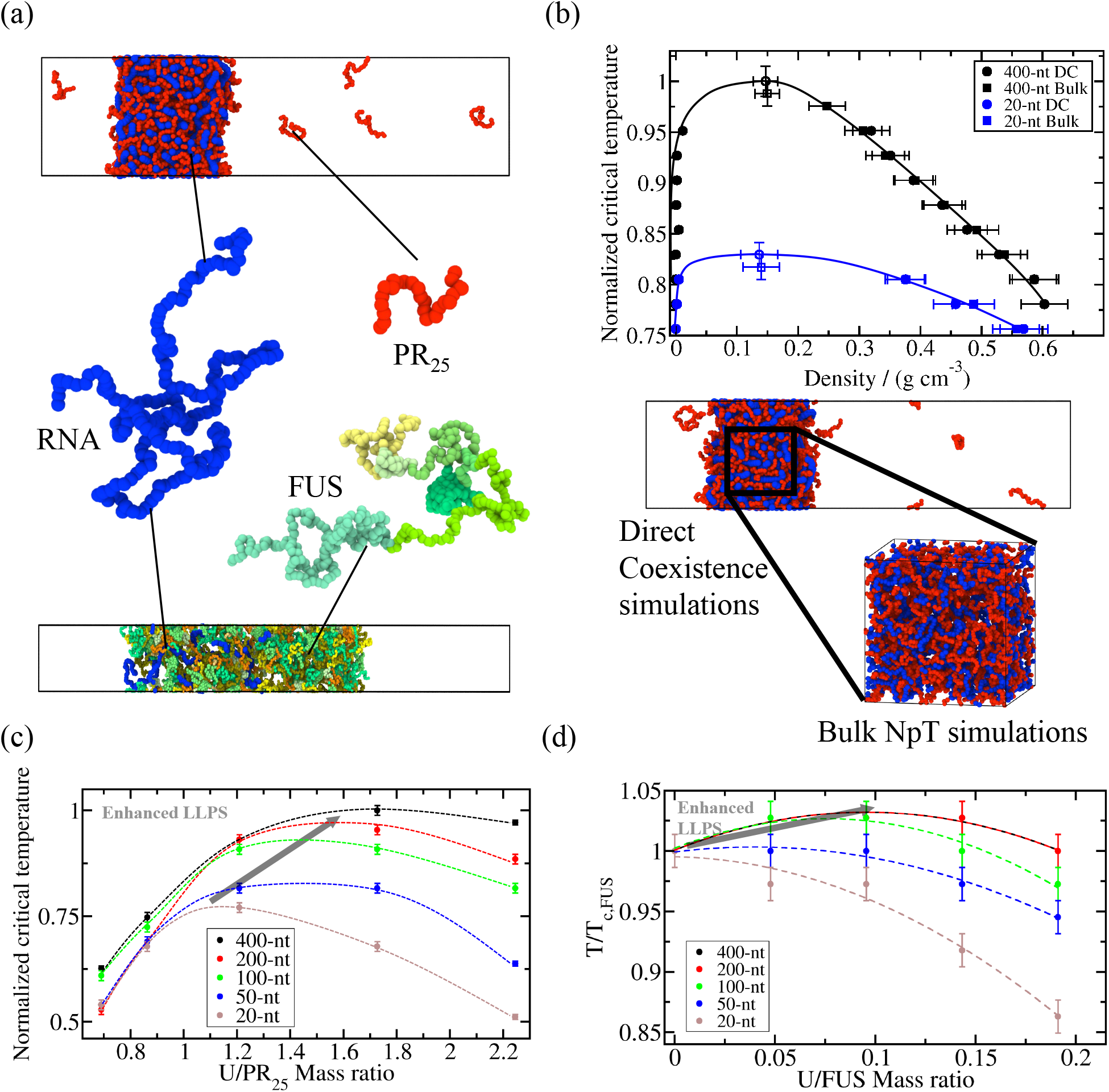
Reentrant phase behaviour driven by RNA is regulated by both concentration and length. (a) Residue-resolution coarse-grained simulations with the Mpipi model [73] to investigate phase-separation of RNA-protein mixtures. Coarse-grained representation of (full-sequence) FUS, PR_25_, and a 400-nt polyU RNA strand using the Mpipi model [73] in which each amino acid or nucleotide is represented by a single bead. Please note that the size of the beads depicted in this panel has been conveniently rescaled for visualization purposes. In FUS protein, beads of different colours indicate distinct protein domains. Direct Coexistence simulations of polyU-PR_25_ (Top) and FUS-polyU (Bottom) are also included. (b) Comparison of the predicted condensate densities as a function of temperature (renormalized by the highest critical temperature) for polyU-PR_25_ mixtures composed by polyU strands of 400-nt (black symbols) and 20-nt (blue symbols) using Direct Coexistence simulations (filled circles) and Bulk *NpT* simulations (filled squares). The estimated critical temperature of each system by both ensembles is depicted by empty symbols of the corresponding shape and colour. Snapshots of a Direct Coexistence simulation and a Bulk *NpT* simulation are included to illustrate the analogy between both ensembles when describing the system condensed phase. (c) Normalized critical temperature of polyU-PR_25_ mixtures as a function of the U/PR_25_ mass ratio for different polyU strand lengths as indicated in the legend. (d) Normalized critical temperature of FUS-polyU mixtures as a function of the U/FUS mass ratio for different polyU strand lengths as indicated in the legend. While in Panel (c) all temperatures have been normalized by the highest T at which phase separation was observed (T= 435K), in Panel (d) all temperatures have been normalized by the critical temperature of pure FUS (T_*c,F US*_=365K). Please note that higher critical temperatures in our model correspond to higher driving forces to undergo LLPS (i.e., lower saturation concentration).

In addition to the residue-resolution Mpipi model, we employ a minimal coarse-grained patchy model (MD-Patchy [64]) to elucidate whether the role of RNA length and concentration in RNA-protein condensates is determined by general molecular features such as valency, binding affinity or the relative RNA/protein length. Within our patchy particle simulations, proteins are described as pseudo hard-sphere (PHS) [91] particles decorated with sticky patches which account for the protein binding sites (modelled through square-well-like potentials [92]), and RNA strands are modelled as fully flexible self-avoiding polymers that can interact attractively with RNA-binding proteins via mid-range non-specific interactions [89]. Specifically, RNA self-avoiding polymers are described by a PHS potential in combination with a Yukawa/Debye-Hückel screened potential (for further details on the model potential and parameters see Section III C, and SM). With our minimal model, each RNA bead accounts for several nucleotides and has the same size as those of the proteins [89]. Moreover, as in the residue-resolution coarse-grained model, an implicit solvent is used. Accordingly, the diluted phase (i.e., the protein-poor liquid phase) and the condensed phase (i.e., the protein-rich liquid phase) are effectively a vapour and a liquid phase, respectively.

To determine the stability of RNA-protein condensates, we evaluate the phase diagrams (in the temperature–density plane) of the different systems by means of Direct Coexistence simulations [93, 94]. Within the Direct Coexistence approach (Fig. 1(a)), the two coexisting phases of a given system are placed in the same simulation box, which is a rectangular box with an elongated side perpendicular to the interfaces—long enough to capture the bulk density of each phase—while the parallel sides are chosen such that proteins and RNA cannot interact with themselves across the periodic boundary conditions [43]. We employ the canonical ensemble (constant number of molecules (*N*), system volume (*V*) and temperature (*T*), or *NV T*). Once Direct Coexistence simulations reach equilibrium, we measure the coexisting densities of both the diluted and condensed phases along the long axis of the box, excluding the fluctuations at the interfaces and keeping the center of mass of the system fixed. By repeating this procedure at different temperatures—until we reach supercritical temperatures, i.e., where no phase separation is observed any longer—we can evaluate phase diagrams (Fig. 1(b)). Finally, to avoid finite system-size effects close to the critical point, we estimate the critical temperature (*T*_*c*_) and density (*ρ*_*c*_) using the law of critical exponents and rectilinear diameters [95] (as shown in Refs. [64, 89]). Fig 1(a) shows a Direct Coexistence simulation with a system composed by PR_25_ and poly-Uridine (polyU) RNA strands of 400 nucleotides (nt) and FUS with polyU strands of the same length at conditions in which both systems undergo LLPS.

Furthermore, to evaluate condensate densities while controlling accurately the concentrations of individual components inside them, we perform additional simulations in the isothermal-isobaric ensemble (*NpT*). In contrast, fixing the desired composition of a multicomponent condensate, e.g. the protein/RNA proportion, within the condensed phase in Direct Coexistence simulations is far from trivial. As shown in Fig. 1(b), *NpT* simulations provide a similar representation of the condensed phase obtained by Direct Coexistence simulations (although avoiding the effects of interfaces). By fixing the system’s pressure to zero, and enabling the volume of the simulation box to isotropically fluctuate, we allow the condensed phase to equilibrate. Stable phase-separated condensates exhibit an equivalent coexisting density as that obtained by Direct Coexistence simulations, whereas unstable systems cannot sustain the condensed phase and tend towards infinitely dilute densities. Within the *NpT* ensemble, condensates are only stable at temperatures where the interactions between biomolecules are sufficiently strong to overcome the entropic cost of forming a percolating liquid network without the need for pressure to be exerted on the simulation box. Finally, once the systems’ densities are equilibrated, the critical temperature of each condensate can be estimated within the interval between the highest temperature at which the system is stable and the lowest temperature at which it is not. A comparison between the condensate coexistence densities and critical temperatures obtained via *NpT vs*. Direct Coexistence simulations for mixtures of polyU-PR_25_ with different strand lengths is shown in Fig. 1(b).

## III. RESULTS AND DISCUSSION

### A. RNA length modulates the RNA concentration-dependent reentrant phase behaviour of RNA-protein condensates

We first investigate how the RNA-concentrationdependent reentrant phase behaviour of RNA-protein condensates is influenced by the length of single-stranded RNA. For this, we perform DC and bulk *NpT* simulations using the residue/nucleotide-resolution Mpipi force field [73], which has been shown to achieve quantitative agreement with experimental phase diagrams of RNA-binding proteins. We compare the behaviour of two phase-separating RNA-binding proteins, FUS (526 residues, sequence in Suplementary Material) and PR_25_, for which phase behaviour is modulated differently by RNA [44]. FUS can form single-component condensates via homotypic interactions, which increase in stability at moderate RNA concentrations [37, 43, 46]. PR_25_ is an arginine-rich peptide, which requires RNA to phase separate via heterotypic RNA–protein interactions at physiological conditions [44, 51]. For each case, we simulate solutions containing tens to hundreds of individual proteins in the presence of disordered single-stranded polyU RNA molecules with five different lengths: 20, 50, 100, 200, and 400 nucleotides. For all RNA lengths, we test different polyU concentrations defined through the U/protein mass ratio, which allows us to quantify the total number of U nucleotides in the mixtures, regardless of whether they are assembled in longer or shorter polyU chains. Importantly, it has been shown that combining RNA-binding proteins and RNA at ratios resulting in electroneutral mixtures enhances the stability of RNA-protein condensates [42, 43]. Thus, here we explore a range of U/protein mass ratios that lie around the electroneutral point. We focus on single-stranded polyU RNA for simplicity and to follow previous works on RNA-protein phase separation [18, 37, 74].

For the polyU-PR_25_ mixtures, the electroneutral point lies at the 1.21 U/protein mass ratio. Thus, we perform simulations for polyU-PR_25_ mixtures spanning the range between 0.78 and 2.24 U/PR_25_ mass ratios. For the electroneutral polyU-PR_25_ system, we first demonstrate that *NpT* simulations quantitatively reproduce the condensed phase coexistence densities found in phase diagrams constructed using standard *NV T* Direct Coexistence simulations [64, 84]. From a set of Direct Coexistence simulations at varying temperatures, we extract the phase diagrams for two polyU-PR_25_ mixtures containing RNA strands of varying lengths (20 and 400 nucleotides) but keeping a constant U concentration (mass ratio of 1.21) regardless of RNA length (Fig. 1(b)). Then, we simulate these systems in the *NpT* ensemble at *p*=0 (using a cubic box). As shown in Fig. 1(b), this approach provides consistent condensate densities to those obtained via Direct Coexistence simulations (as long as the density of the dilute phase is very low). As our Direct Coexistence simulations indicate, this assumption is reasonable for most of the temperatures with densities of the dilute phase being of the order 1×10^−3^ g/cm^3^ (please note that the solvent is implicitly considered within the force field). Since with *NpT* simulations the density of the dilute phase cannot be measured, the value for the critical temperature cannot be calculated using the law of critical exponents and rectilinear diameters [95], as is the case in the Direct Coexistence simulations. However, an interval at which the critical temperature falls can be estimated. Figure 1(b) shows that the values of the critical temperatures estimated from bulk *NpT* simulations (as the mid temperature of the computed interval; described in Section II) lie within the uncertainty of the critical temperatures evaluated from Direct Coexistence simulations and the law of critical exponents and rectilinear diameters.

Having established that *NpT* simulations can provide robust estimates of the critical temperature, we move forward with the *NpT* ensemble to investigate the interplay between RNA concentration and the effects of RNA length on condensate stability. To consider the effect of RNA length independently, we keep the total amount of U nucleotides in the mixtures fixed, and assemble them in polyU chains of different lengths. When the U/PR_25_ mass ratio is kept constant, we observe a monotonic increase in the critical temperature of the condensates as the RNA length increases. For example, for polyU-PR_25_ mixtures at ratios satisfying the electroneutral point (i.e., 1.21 polyU/PR_25_ mass ratio), there is a 20% enhancement in the critical temperature when the RNA chain length increases from 20 to 400 nucleotides (Figure 1(c)). When we next fix the RNA chain length, and investigate the impact of RNA concentration on the stability of polyU-PR_25_ condensates, for all the RNA lengths we study, we confirm that PR_25_ exhibits the well-known RNA-concentration dependent reentrant behaviour of RNA-binding proteins discovered experimentally [22, 37, 46]. That is, the stability of polyU-PR_25_ condensates—quantified by the values of the critical temperature—gradually increases as the RNA concentration goes from low to moderate (up to approximately the electroneutral point, at 1.21 mass ratio), then it reaches a maximum value, and finally decreases as the RNA concentration increases even further. Remarkably, looking at the combined effects of RNA length and concentration reveals that RNA length significantly modulates such reentrant behaviour. Specifically, increasing RNA length sensitively raises the maximum value that the critical temperature of the mixture reaches (i.e. how much the condensate stability can be boosted by RNA), and the maximum concentration of U nucleotides that the condensate can incorporate before beginning to become unstable (i.e., when the RNA chains are longer, more nucleotides can form part of the condensate before it begins to dissolve). Thus, the most stable polyU-PR_25_ condensates are formed by the longest RNA chains we study, and, unexpectedly, contain U/PR_25_ concentrations above the electroneutral point (Fig. 1(c)).

We next investigate whether this behaviour also holds for proteins that are able to undergo phase separation on their own, i.e., via homotypic protein–protein interactions [65, 96]. For this, we focus on the protein FUS, and test the impact of adding polyU of varying lengths and at different concentrations. Our simulations contain 4 copies of FUS (full-sequence; see SM) and polyU chains of either 20-nt, 50-nt, 100-nt, 200-nt or 400-nt in length. For all the different polyU lengths, we prepare mixtures at concentrations spanning the range of U/FUS mass ratio from 0 to 0.19. First, when we look at mixtures containing RNA at the concentration corresponding to the electroneutral point (mass ratio of 0.049), we confirm that performing *NpT* bulk simulations reproduces the length-dependent increase in critical temperature identified previously via Direct Coexistence simulations [44]. Specifically, when we mix FUS with RNA of either 100-nt, 200-nt or 400-nt at the electroneutral ratio (0.049 U/FUS mass ratio), we see a marginal increase in critical temperature—3% with respect to the value for pure FUS system (Figure 1(d))—in agreement with our previous Direct Coexistence results [44]. Adding 20-nt polyU to FUS, the shortest polyU molecules we study, hinders phase separation, with the effect being amplified at higher polyU concentrations. This occurs because 20-nt polyU is too short to bridge FUS molecules and enhance the conectivity of the condensed liquid, as shown previously [43, 44]. For RNA lengths of 50-nt, 100-nt, 200-nt and 400-nt, the RNA concentration-dependent reentrant phase behaviour of FUS is observed. That is, gradually increasing the concentration of polyU up to a given threshold (in our case a U/FUS mass ratio of around ∼ 0.1) increases the critical temperature, whilst adding polyU at concentrations beyond such threshold reduces the critical temperature. The observation of peak critical temperatures for FUS condensates containing 50-nt polyU at polyU concentrations surpassing the electroneutral point (i.e., ∼ 0.1), is consistent with *in vitro* studies [20]. As in PR_25_/polyU mixtures, we find that the maximum enhancement in phase separation occurs at higher polyU/protein mass ratios for longer chain polyU systems. Overall, when polyU length is increased from 20 to 400 nucleotides, the mass ratio at which the system displays the highest critical temperature increases from 0 (no RNA) to 0.12 (Figure 1(d)). Nevertheless, we observe that the RNA length-dependent effects for FUS/polyU mixtures are significantly smaller than for polyU-PR_25_ mixtures (see Figs. 1(c) and 1(d)).

Our results for FUS are consistent with a number of *in vitro* studies which identified similar patterns of RNA concentration-dependent reentrant phase behaviour [41, 46]. However, that low levels of RNA promote phase separation, whilst higher RNA levels promote dissolution contrasts with the observation that some condensates are able to form in regions of the cell with very high levels of RNA (up to RNA/protein mass ratios of ∼40) [97]. Moreover, despite RNA being at high concentrations, it has been shown to play a key role in initiating phase separation of FUS by nucleating protein condensates [20, 46, 97]. For example, RNA plays a prominent role in triggering P-bodies formation, which occurs as a result of a strong increase in mRNA concentration [97, 98]. Also, FUS-containing paraspeckles in the nucleus are thought to be nucleated by the long non-coding RNA Neat1, despite the high background RNA concentration [20, 46]. Even though *in vitro* studies have not explicitly revealed how the length of RNA defines the maximum RNA concentration that a condensate rich in RNA-binding proteins can take before its stability begins to decrease, there is evidence which indirectly supports this theory: a number of *in vitro* and *in silico* studies have identified that minimum RNA lengths are needed to promote phase separation of RNA-binding proteins [43, 99] and others have found that shorter RNA molecules are more potent promoters of condensate dissolution [46], in agreement with our results from Figure 1(d). Here, we show directly that increasing the length of the nucleic acid chain increases the capacity of the RNA to promote phase separation and to be incorporated into the condesate at higher RNA concentrations than their shorter counterparts. Therefore, previous findings which have shown that high concentrations of RNA promote condensate dissolution ought to be contextualised with information on the length of the studied nucleic acids. As demonstrated here, it is possible for phase separation to be favoured at high RNA concentrations if the RNA chains are long enough to bind multiple proteins simultaneously. Assembling nucleotides in longer RNA chains has the additional advantage of decreasing the electrostatic repulsion among phosphates of separate nucleotides, due to the presence of more covalent bonds.

Since our results support the hypothesis that longer RNA molecules are more powerful enhancers of phase separation due to their ability to increase the connectivity of the condensed liquid [66, 96], we next investigate this behaviour by examining the stability of condensates containing both long and short polyU chains (Fig. 2(a)). To do so, we simulate mixtures of PR_25_ in the presence of polyU strands of two different lengths, 50-nt and 400-nt, such that each length represents half of the total polyU mass ratio concentration in each system. For these mixed RNA length systems, we explore polyU/PR_25_ mass ratios ranging from 0.76 to 2.1. We find that mixed RNA length systems display critical temperatures in between those of the pure short and long RNA chain systems, and with the maximum enhancement of critical temperature also occurring at intermediate polyU/PR_25_ mass ratios between that of the pure short and pure long RNA systems (Figure 2(b)). We also study FUS-polyU mixtures of 50-nt and 400-nt polyU strands (such that each length represents half of the total polyU mass ratio concentration) spanning mass ratios from 0 to 0.196. Similar to the results for PR_25_, the mixed length systems in FUS present intermediate critical temperatures compared to the pure long-chain and short-chain RNA systems, as well as patterns of reentrant phase behaviour in which the maximum critical temperature peaks at an intermediate polyU/FUS mass ratio between that of the pure long-chain and short-chain RNA systems (Figure 2(c)).

**FIG. 2:**
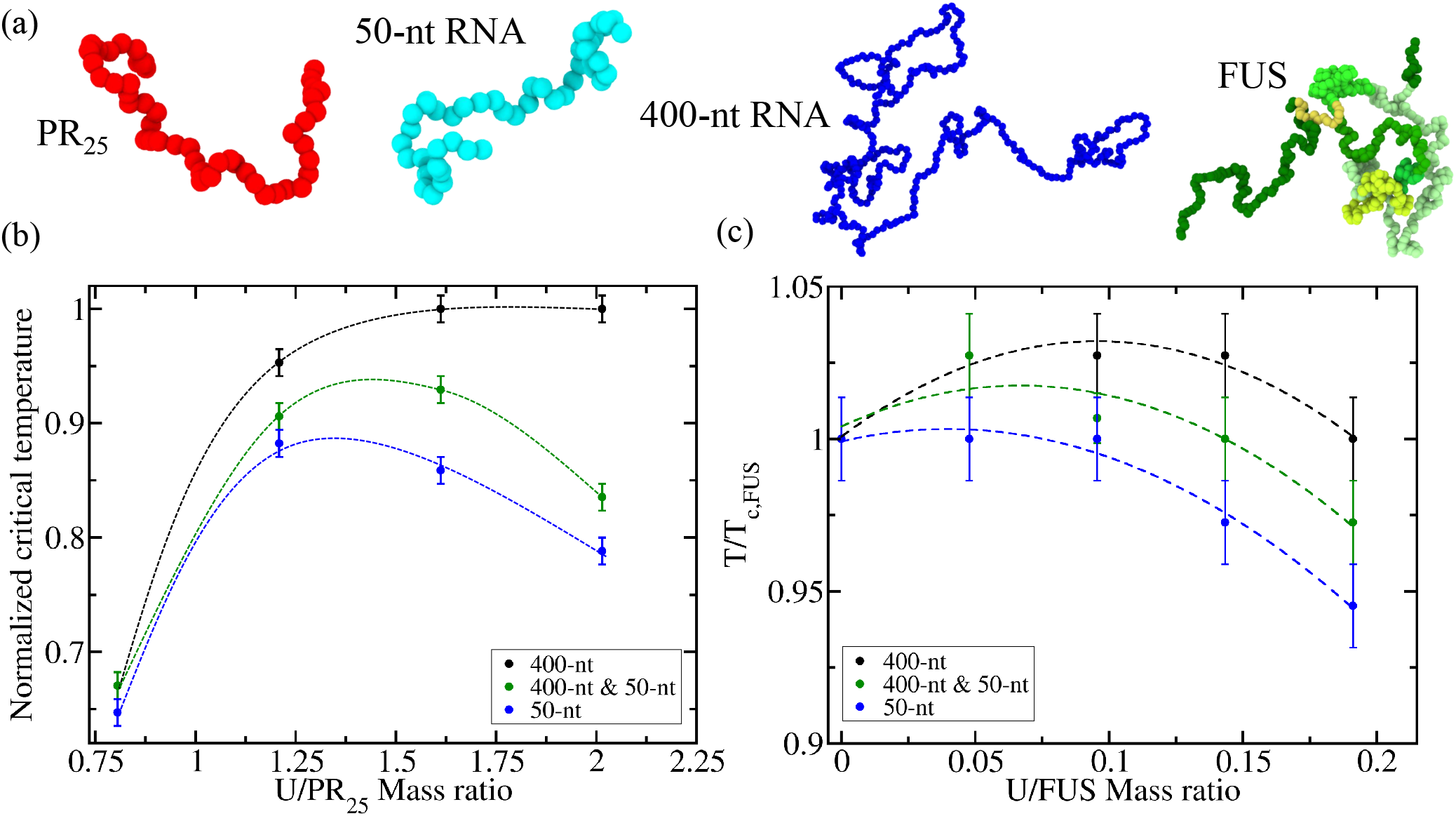
RNA-driven reentrant phase behaviour of FUS and PR_25_ polyU systems including mixtures with different RNA strand lengths. (a) Representation of PR_25_, FUS, and two RNA strands of 50 and 400-nucleotide each following the same colour code and considerations discussed in Fig. 1(a). Please note that the size of the beads depicted in this panel has been conveniently rescaled for visualization purposes. (b) Normalized critical temperature of polyU-PR_25_ mixtures as a function of the U/PR_25_ mass ratio for different polyU strand lengths as indicated in the legend. (c) Normalized critical temperature of FUS-polyU mixtures as a function of the U/FUS mass ratio for different polyU strand lengths as indicated in the legend. For the systems with mixed polyU lengths, each length represents half of the total polyU concentration. While in Panel (b) all temperatures have been normalized by the highest T at which phase separation was observed (T=425K), in Panel (c) all temperatures have been normalized by the critical temperature of pure FUS (T_*c,F US*_).

Our findings reveal that mixed length polyU systems are generally more stable than short-chain polyU systems; further emphasising the ability of long polyU strands to show a stabilising effect on condensates up to higher concentrations, given that we observe this behaviour even when shorter polyU molecules are also present. Our observations are consistent with *in vitro* FUS/RNA phase separation assays, reporting that the lncRNA Neat1 (short isoform, length=3.7kb) was able to drive reappearance of FUS droplets which had previously been solubilised with tRNA (70-100 nucleotides [100]) [46].

### B. RNA length and concentration determine the internal organisation of molecules in RNA-protein condensates and the properties of their interfaces

To rationalise the interaction of RNA length and concentration in modulating the stability of RNA-protein condensates, we now characterise the organisation of the different molecules inside the RNA-protein condensates by quantifying the densities of polyU *versus* proteins across the condensate. These density profiles reveal that there are considerable differences in the distribution of proteins *versus* polyU depending on the concentration of RNA. Protein-rich systems (i.e., low polyU/PR_25_ mass ratios) form condensates with a surface coated by PR_25_ peptides (Fig. 3(a-b)), whereas the surfaces of polyU-rich droplets (i.e., high polyU/PR_25_ mass ratios) are mostly coated by polyU chains (Fig. 3(c-d)).

**FIG. 3:**
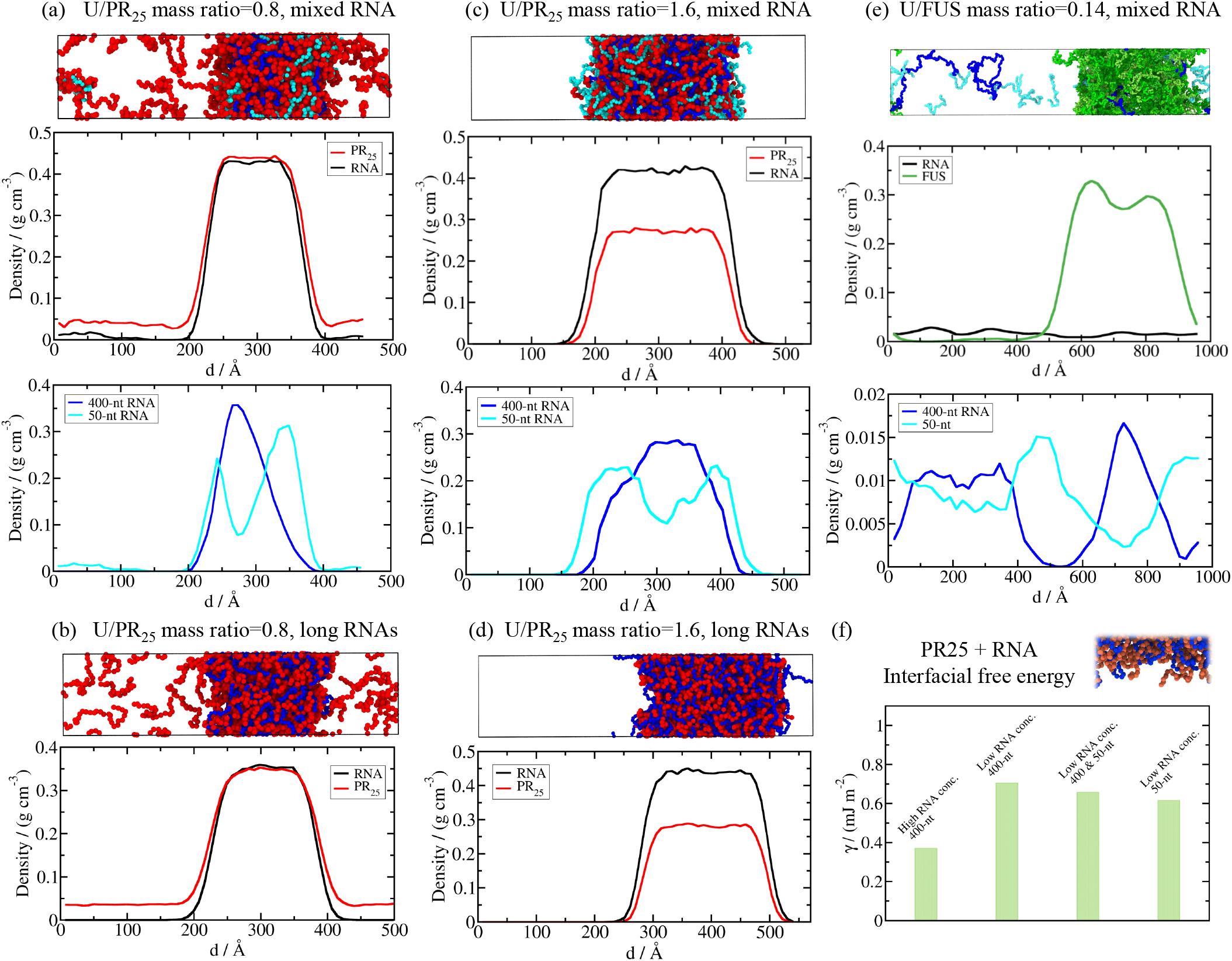
Structural condensate organization of RNA-binding proteins in the presence of long and short RNA strands. (a) PolyU-PR_25_ mixture with a 0.8 U/PR_25_ mass ratio where polyU strands are 50-nt and 400-nt long (each length contributing half to the total polyU concentration). Top: Representative snapshot of a Direct Coexistence simulation of the system, where PR_25_ molecules are coloured in red and long and short RNAs are depicted in blue and cyan respectively. Middle: Density profile of PR_25_ (red) and polyU RNA (black) along the long axis of the simulation box. Bottom: RNA density profile decomposed in 400-nt (blue) and 50-nt (cyan) polyU chains. (b) The same as in Panel (a), but for a polyU-PR_25_ mixture with a U/PR_25_ mass ratio of 0.8 and polyU strands of 400-nt only. Note that we only show one density profile since here all RNAs are of equal length. (c) The same as in Panel (a), but for a polyU-PR_25_ mixture with a U/PR_25_ mass ratio of 1.6, where polyU strands are also 50 and 400-nt long. (d) The same as in Panel (b), but for a polyU-PR_25_ mixture with a U/PR_25_ mass ratio of 1.6 and polyU strands of 400-nt only. The temperature of systems shown in Panels (a)-(d) was 0.9 with respect to their corresponding critical temperature. (e) FUS-RNA mixture at T/T_*c,F US*_=0.98 and with a U/FUS mass ratio of 0.14, where polyU RNA strands are 50-nt and 400-nt long (each length contributing half to the total polyU concentration). Top: Representative snapshot of a Direct Coexistence simulation. Middle: Density profile of FUS (green) and RNA (black) along the long axis of the simulation box. Bottom: RNA density profile decomposed in 400-nt (blue) and 50-nt (cyan) polyU chains. (f) Surface tension for different polyU-PR_25_ mixtures, all of them at a temperature of 0.85 with respect to its corresponding critical temperature for the system indicated in the legend. The high RNA concentration corresponds to a U/PR_25_ mass ratio of 1.6, while the low RNA concentration corresponds to a U/PR_25_ mass ratio of 0.8.

In polyU-PR_25_ condensates combining RNAs of different lengths, we find that the RNAs exhibit a striking spatially heterogenous distribution. The longer 400-nt polyU molecules accumulate in the condensate core, while the shorter 50-nt polyU molecules concentrate preferentially towards the condensate surface (Figs. 3(a) and 3(c)). Simulations approximating proteins as patchy colloids have revealed the same heterogenous organisation of high and low valency proteins within multicomponent condensates. By looking at the problem from a condensed matter perspective, the simulations revealed that burying high valency proteins in the centre, and exposing low valency species to the interface, maximises the enthalpic gain for condensate formation (most bonds of the higher valency molecules are satisfied at the condensate core) and reduces the interfacial free energy at the droplet interface because the lower valency molecules act as surfactants [63, 66, 101]. In Figure 3(e), we also show that polyU-FUS condensates of mixed RNA length (with equal concentration of 50-nt and 400-nt strands) display multilayered RNA organisation, preferentially locating short polyU chains at the interface and long RNA strands in the core. Such structural organisation maximizes at the same time the condensate liquid network connectivity and minimizes the interfacial penalty.

Since distinct molecules at the condensate interface can translate into significantly different interfacial properties, such as interfacial free energies [22, 43], coalescence fusion rates [102], size-conservation [6, 63], or uneven molecule exchange rates [66], we now calculate the interfacial free energy (*γ*; see SM for further details on these calculations). We focus on PR_25_ condensates because converging the value of *γ* for FUS-based droplets is computationally unfeasible due to the size of FUS (526 amino acids). We start by computing the interfacial free energy for two types of 400-nt polyU-PR_25_ condensates: with a PR_25_-rich interface (i.e., low U/PR_25_ mass ratio of 0.8) and with a polyU-rich interface (i.e., high U/PR_25_ mass ratio of 1.6), as shown in Figure 3(f). To faciliate the comparison, in both cases we simulate the systems at the same temperature (*T/T*_*c*_=0.85). Strikingly, we find that the condensate with a polyU-rich interface presents a much lower interfacial free energy (almost half) than that with a PR_25_-rich interface. Such an unexpected finding explains how a high concentration of polyU, beyond the electroneutral point, can boost the stability of condensates: by incorporating a large concentration of RNA, condensates experience a trade-off between the destabilising effect of a decreased enthalpic gain due to the larger electrostatic repulsion among equally charged nucleotides, and the stabilising effect of the decrease in the condensate energetic penalty of forming an interface when it is coated with polyU. Our results, therefore, reveal that in condensates with an excess of polyU, polyU behaves like a surfactant.

To analyse the impact of RNA length, we now calculate the interfacial free energy of two different condensates containing the shorter 50-nt polyU RNA: a PR_25_, 50-nt and 400-nt polyU condensate, and a PR_25_ and 50-nt polyU condensate. We estimate the interfacial free energy at the same temperature used above (*T/T*_*c*_=0.85) and at the lower U/PR_25_ mass ratio of 0.8 (*T/T*_*c*_=0.85), since imposing a high RNA concentration with short polyU molecules within the condensed phase in a Direct Coexistence simulation is not feasible. We observe that increasingly adding 50-nt polyU molecules progressively decreases the interfacial free energy of the condensate, with respect to the value of the 400-nt polyU-PR_25_ condensate (Fig. 3(f)). This observation reinforces the idea that in RNA-protein condensates that contain RNAs of different lengths, positioning the shorter RNAs species at the interface reduces the surface tension as such shorter RNAs act as better surfactants than longer ones [103]. Hence, the advantage of mixed length RNA condensates exhibiting multilayered organization results from them presenting similar low surface tensions as those only formed by short RNA strands, while showing considerably higher stability due to long RNA strands increasing the enthalpic gain for condensate formation by contributing more connections to the liquid network connectivity together with PR_25_.

Multilayered condensates such as those found in Figs. 3(a), (c) and (e) for PR_25_ and FUS-polyU mixtures respectively can be found across the cell and include FUS-containing paraspeckles [9], stress granules [30] and the nucleolus [6]. Indeed, condensate structure and organisation has important implications for the behaviour of the various components, with those located in the core exhibiting slower exchange rates compared to molecules in the outer shell [66, 96]. Experiments and simulations reveal that longer RNA strands lead to higher viscosities in RNA-protein condensates [18–20, 43, 104]. Thus, in mixed RNA condensates, a stable core containing long polyU strands is expected to have a higher viscosity, while an outer shell with short polyU strands a lower viscosity [43]. Gelification of RNA-protein condensates via fibrillation, with potential pathological implications [21, 47, 105], has been shown to be seeded at the interface due to a local increase of protein density at the surface [106]; hence, incorporating short RNAs into RNA-protein condensates might contribute to preventing their maturation because it decreases the probability of high density fluctuations at the interface.

### C. Self-avoiding polymers trigger concentration-dependent reentrant phase behaviour of colloidal patchy-particle condensates modulated by polymer length

To investigate whether the observed patterns of RNA-length-and-concentration-dependent phase behaviour found in RNA-protein condensates rely on general molecular features such as protein valency, binding affinity, or the relative polymer size/length between proteins and RNA, we employ a minimal CG model of colloidal patchy particles with self-avoiding polymer chains to mimic proteins and RNA respectively [89]. Iterations of this minimal coarse-grained model have been previously used to investigate critical factors in LLPS such as surface tension, droplet size conservation, or condensate substructure [44, 66]. The aim of our minimal simulations here is to assess, beyond protein sequence and specific molecular features, the thermodynamic parameters that explain the general differences between the impact of RNA length and concentration on homotypic phase separation *versus* RNA–protein heterotypic complex coacervation.

We start by computing the phase diagrams of two different types of colloidal patchy particles in the presence of different concentrations and lengths of a self-avoiding polymer that mimics RNA. The first type are patchy particles decorated with 3-binding sites in a planar arrangement separated by 120 degrees angles (Fig. 4(a); see SM for further details on the model). Like FUS, these colloidal particles—referred to henceforth as scaffold proteins—are able to phase separate on their own via homotypic interactions below a reduced temperature of T*=0.09 [89] (see details on reduced units in the SM). On the other hand, the second type of patchy colloids possess 2-binding sites in a polar arrangement, which by construction can only form linear chains and not 3-dimensional percolated networks that sustain phase-separation [62, 64, 65] (Fig. 4(a)). Like PR_25_, 2-binding site colloidal particles—referred to henceforth as cognate proteins—cannot phase separate on their own [44]. We perform Direct Coexistence simulations of both scaffold and cognate proteins for different polymer-bead/protein ratios using polymer chains of 10, 20, 50 and 100 beads such that the polymer-bead/protein ratio is defined in terms of the number of polymer and protein beads, ranging from 0.2 to 1.

**FIG. 4:**
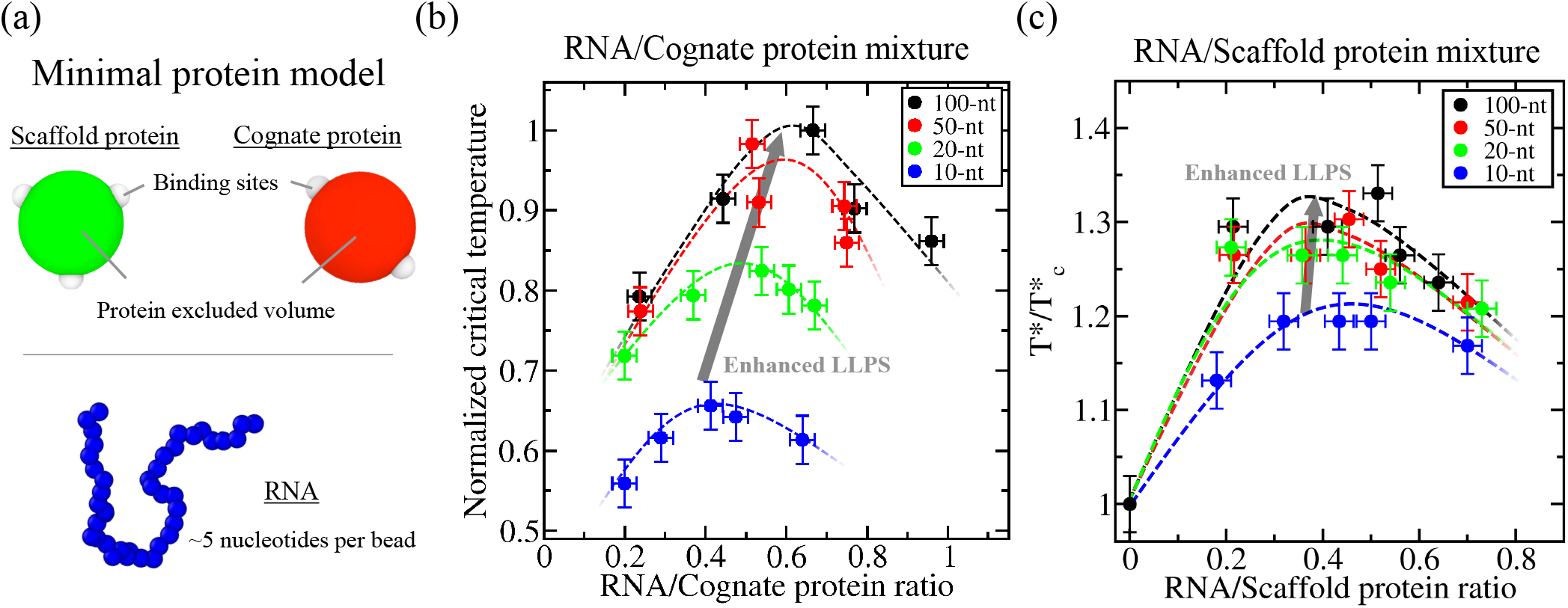
Minimal coarse-grained model for protein LLPS. (a) Green and red spheres represent the excluded volume of scaffold and cognate proteins respectively, while gray patches represent the binding sites of the proteins. Two different proteins are modeled: scaffold proteins, with 3 binding sites in a planar equidistant arrangement, and cognate proteins, with 2 binding sites in a polar arrangement. Blue spherical beads account for ∼5 nucleotides each in the RNA model. Please note that for visualization purposes, the size of the RNA beads has been scaled down. For further technical details on the model, please see the Supplementary Material. (b) Normalized critical temperature of RNA/cognate protein mixtures as a function of the RNA/cognate protein ratio for different RNA strand lengths as indicated in the legend. (c) Normalized critical temperature of RNA/scaffold protein mixtures as a function of the RNA/scaffold protein ratio for different RNA strand lengths as indicated in the legend. While in Panel (b) all temperatures have been normalized by the highest T (T*=0.11) at which phase separation was observed, in Panel (c) all temperatures have been normalized by the critical temperature of the scaffold protein in absence of RNA 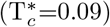.

First, by focusing on a given polymer-bead/protein ratio (i.e., 0.4), we capture the length-dependent enhancement of LLPS reported in our sequence-dependent CG simulations of PR_25_ and FUS (Figure 1) where systems containing longer RNA strands can phase-separate up to higher critical temperatures (Fig. 4(b-c)). For cognate colloidal particles, increasing the ‘RNA’ chain length from 10 to 100 beads at an RNA/protein ratio of 0.4 leads to a 50% increase in critical temperature (Fig. 4(b)). In contrast, the enhancement in critical temperature for the same polymer/protein concentration with scaffold proteins is just 8% (Fig. 4(c)). Clearly, scaffolds show much more subtle dependency on RNA length compared to cognate proteins, in full agreement with our previous sequence-dependent simulations shown in Figs. 1 and 2.

Secondly, by gradually varying the polymer-bead/protein ratio (from 0.2 to 1; Fig. 4(b-c)), patterns of concentration-dependent reentrant phase behaviour for both types of proteins and distinct RNA polymer lengths can be observed. A moderate increase in the polymer-bead/protein ratio leads to an enhancement in the critical temperature, while any further increase in self-avoiding polymer levels results in LLPS inhibition (i.e., lower critical temperatures). For the cognate protein mixtures, we find a significant shift in the ratio which gives the highest critical temperature as a function of polymer length; consistent with our results for PR_25_ with polyU (Fig. 1(c)). On the other hand, for the scaffold protein mixtures, the shift in the maximum critical temperature towards higher polymer/protein ratios with length is more modest; also in qualitative agreement with our polyU-FUS simulations shown in Fig. 1(d). Remarkably, by means of the colloidal patchy particle model, we can explore mixtures with polymer RNA chains much longer than the size of the proteins. We discover that while condensates with RNA lengths of 50 or 100 beads are stable up to higher critical temperatures compared to those of shorter lengths (i.e., 20 beads), for all polymer-bead/protein ratios and for both scaffold and cognate proteins, beyond a certain length at which RNA greatly exceeds the size of the proteins (e.g., 50 times longer than the protein size), the effect of RNA length on LLPS becomes extremely mild (Fig. 4(b-c); red and black curves). These results also support the notion that the longer RNA strands promote condensate stability by increasing the connectivity among proteins within the condensate network [43, 44]. Moreover, since even the shortest polymer length added to the scaffold/cognate mixtures meets this criterion, we do not observe the disruptive influence on LLPS of extremely short RNA chains (i.e., 20-nt) with FUS (526 amino acids) observed in Fig. 1(d).

Our results are consistent with the experimental observation that longer RNA strands present weaker dissociation constants with N-RRM1-2 domains of TDP-43 (which, like PR_25_, cannot phase separate on their own at physiological conditions) than 3-fold shorter RNA strands [33]. It has also been shown that length and charge segregation in the IDR domain of VRN1-like proteins has a critical impact on modulating the DNA-induced VRN1 phase separation, where liquidlike, gel-like or no phase-separation behaviour can be switched depending on the IDR length and the presence of neutral *vs*. highly charged domains [107]. Overall, our patchy particle results presented here, highlight that the RNA length-and-concentration-dependent reentrant phase behaviour observed for both homotypic and heterotypic phase-separating proteins is a general property of soft-matter systems, like biomolecules. Therefore, general features of biomolecules, such as their valency, topology, binding affinity and relative length or size, is what ultimately dictates the intricate phase behaviour of multicomponent biomolecular condensates.

## IV. CONCLUDING REMARKS

In this study, using a multiscale modelling approach we reveal how RNA length can tune concentration-dependent reentrant phase behaviour of RNA-protein condensates. Drawing together the results from our sequence-dependent model and patchy particle simulations, we demonstrate that long RNA polymers act as enhancers of phase separation. Not only do longer RNA strands enable phase separation up to higher critical temperatures, they also facilitate phase separation up to higher RNA/protein ratios. We show that this is a physical feature of long polymers, since consistent mass ratios and net charges are used for all our RNA/polymer systems with different lengths. Our finding that longer chain RNA molecules increase the capacity of RNA to raise the critical solution temperature to higher values and that the corresponding condensates can incorporate a higher proportion of RNA nucleotides had not been directly demonstrated previously. However, these results are consistent with the wide body of work demonstrating the stabilising role of molecules with an increasing number of repeating chemical building blocks, or higher valencies, on biomolecular condensates (e.g., longer RNAs [44], DNAs [108, 109], or polySUMO/polySIM [96], and IDPs with more stickers [35, 56, 110]).

Additionally, our finding that long RNA tends to localise to the condensate’s core whereas shorter strands accumulate at the droplet surface provides further support for the theory that the stabilising and enabling effects of long RNA chains comes, in part, from their ability to act as nucleators [46] and scaffolds [44] in phase-separated condensates. While long RNA stabilise condensates by increasing the density of molecular connections of the liquid network, short RNAs act as better surfactants that increase condensate stability by reducing their interfacial free energy [63]. Moreover, the influence of RNA length on condensate organisation and viscoelastic properties has important implications on the dynamics of the different components and the mechanisms explaining liquid-to-solid transitions, with potential implications in neurodegenerative disorders [18, 19, 21, 47, 102].

Considering the significant interplay between RNA length and concentration might be relevant to rationalise why some biomolecular condensates, such as stress granules or paraspeckles, can be nucleated by long RNA strands [20, 46, 97] despite being in RNA-rich environments. According to our results, such behaviour might be facilitated by RNAs serving as scaffolds and surfactants of RNA-protein condensates at high concentration. In particular, while high RNA-concentrations are expected to inhibit phase separation [53], a diverse population of shorter and longer RNAs can yield stable condensates by incorporating the longer RNAs at concentrations beyond the electroneutral point at the condensate core—to enhance condensate connectivity— and the shorter RNAs at the condensate interface—to reduce the interfacial free energy. The observation that RNA length has a similar impact on the concentrationdependent reentrant phase behaviour for both FUS and PR_25_ condensates, although quantitatively shifting the maximum critical temperature by substantially different extents, is also significant and further emphasises that this observation results from the physical properties of long RNA chains rather than merely on the type of interaction driving phase separation.

Our results also provide evidence for the hypothesis by Henninger *et al*. [53] that RNA length and concentration may act in concert to regulate the formation of intranuclear condensates. Specifically, the hypothesis that formation and dissolution of transcriptional condensates is regulated by RNA through a negative feedback loop [53] could be explained by RNA-concentrationdependent reentrant phase behaviour alone. However, our findings suggesting that longer chain RNAs, which are produced during a transcriptional burst, stabilise condensates up to higher RNA concentrations may provide robustness to the feedback loop. As higher nucleotide concentrations would need to be reached for condensate dissolution, the transcriptional condensates would remain intact until sufficient levels of longer chain mRNA have been synthesised.

Taken together, our results demonstrate some of the many ways in which RNA act as a key regulator of phase separation in the cell and adds to the growing consensus considering its role as essential for developing a robust understanding of the regulation and dysregulation of biomolecular condensates [21, 111]. Several causes of condensate damage in neurodegenerative disorders, such as disease associated mutations [112, 113], do not alter the fundamental changes in phase behaviour which differentiate liquid-like condensates from solid-like aggregates [102], but act to increase the likelihood of these changes occurring [105]. In other words, the properties of condensates containing mutated proteins might not be discernibly different from those containing wild type proteins other than in terms of the timescale of their maturation to solid-like aggregates. This is a strong indication that the focus of research into potential treatments for condensate-associated diseases should shift towards more fundamental and general mechanisms of biomolecular condensate regulation [114–117]. In that sense, the growing body of evidence that RNA plays a major role in driving changes of condensate phase behaviour, to which this work contributes, positions RNA-protein liquid-liquid phase separation as an area of research with great potential.

## ACKNOWLEDGEMENTS

This project has received funding from the Oppenheimer Research Fellowship of the University of Cambridge. I. S.-B. acknowledges funding from the Oppenheimer Fellowship, Derek Brewer scholarship of Emmanuel College and EPSRC Doctoral Training Programme studentship, number EP/T517847/1. J. R. E. also acknowledges funding from the Roger Ekins Research Fellowship of Emmanuel College. R.C.-G. acknowledges funding from the European Reseach Council (ERC) under the European Union’s Seventh Framework Programme (FP7/2007-2013) through the ERC grant PhysProt (agreement no. 337969). This work has been performed using resources provided by the Cambridge Tier-2 system operated by the University of Cambridge Research Computing Service (http://www.hpc.cam.ac.uk) funded by EPSRC Tier-2 capital grant EP/P020259/1.

## SUPPLEMENTARY INFORMATION

Please see the Supplementary Material (SM) for further details on the models, simulation details, and computational methods employed in this work.

