## Supplementary Material for "Surfactants or scaffolds? RNAs of different lengths exhibit heterogeneous distributions and play diverse roles in RNA-protein condensates"

### I. MODELS AND SIMULATION DETAILS

#### A. Mpipi Model

For our simulations of *fused in sarcoma* (FUS) (see sequence below) and $PR_{25}$ with polyU RNA, we employ the high-resolution sequence-dependent coarse-grained model Mpipi [1], which describes almost quantitatively the temperature-dependent LLPS phase behaviour of different protein condensates such as that of FUS. In addition, this model correctly predicts the multiphase behaviour of the polyR/polyK/polyU system, and recapitulates experimental LLPS trends for sequence mutations on FUS, DDX4 NTD and LAF-1 RRG domain variants [1]. Within this force field, electrostatic interactions are modelled through a Coulombic term with Debye–Huckel electrostatic screening [2], given by the sum over all particle-particle $(i, j)$ interactions as:

$$E_{elec} = \sum_{i,j} \frac{q_i q_j}{4\pi\epsilon_r\epsilon_0 r_{ij}} exp(-\kappa r_{ij}) \tag{1}$$

where $q$ is the charge (being -0.75$e$ for the different nucleotides: A, C, G, U; and +0.75$e$ for amino acids such as R and K, +0.375$e$ for H, and -0.75$e$ for D and E residues), $\epsilon_r = 80$ is the relative dielectric constant of water, $\epsilon_0$ is the electric constant, $\kappa^{-1} = 795$ pm is the Debye screening length, and $r_{ij}$ is the distance separating particles $i$ and $j$. For these interactions, a Coulomb cut-off of 3.5 nm is employed. The rest of non-bonded interactions (i.e., hydrophobic, cation-$\pi$ or $\pi$-$\pi$) between distinct protein/RNA beads are modelled via the Wang–Frenkel potential [3]:

$$E_{WF} = \sum_{i,j} \epsilon_{ij}\alpha_{ij} \left[\left(\frac{\sigma_{ij}}{r_{ij}}\right)^{2\mu_{ij}} - 1\right]\left[\left(\frac{\sigma_{ij}}{r_{ij}}\right)^{2\mu_{ij}} - 1\right]^{2\nu_{ij}} \tag{2}$$

where

$$\alpha_{ij} = 2\nu_{ij}\left(\frac{R_{ij}}{\sigma_{ij}}\right)^{2\mu_{ij}}\left[\frac{2\nu_{ij}+1}{2\nu_{ij}\left[\left(\frac{R_{ij}}{\sigma_{ij}}\right)^{2\mu_{ij}} - 1\right]}\right]^{2\nu_{ij}+1} \tag{3}$$

representing $\sigma_{ij}$ the molecular diameter of each residue/nucleotide and $\epsilon_{ij}$ the interaction strength between distinct amino acids and nucleotides ($i$ and $j$). While $\sigma_{ij}$, $\epsilon_{ij}$ and $\mu_{ij}$ are parameters specified for each pair of interactions (see Ref. [1]), $\nu_{ij}$ and $R_{ij}$ are constant model parameters set to $\nu_{ij} = 1$ and $R_{ij} = 3\sigma_{ij}$ for every interaction. Finally, bond energy is computed with an harmonic bond potential of the following form:

$$E_{bond} = \sum_{b} \frac{1}{2}k(r_b - r_0) \tag{4}$$

where $b$ is the total number of bonds, $r_b$ is the bond distance, $k$=8.03 $\mathrm{Jmol^{-1}pm^{-2}}$ is the spring constant and $r_0$ is the bond reference position, set to 381 pm and 500 pm for protein and RNA bonds respectively. For further details on this force field and the full list of the model parameters please see Ref. [1].

#### B. FUS Sequence and PDB of the structured domains

Full-FUS sequence

MASNDYTQQATQSYGAYPTQPGQGYSQQSSQPYGQQSYSGYSQSTDTSGYGQSSYSSYGQSQNTGYGTQSTPQGYGSTGGYGSS
QSSQSSYGQQSSYPGYGQQPAPSSTSGSYGSSSQSSSYGQPQSGSYSQQPSYGGQQQSYGQQQSYNPPQGYGQQNQYNSSSGGGG
GGGGGGNYGQDQSSMSSGGGSGGGYGNQDQSGGGGSGGYGQQDRGGRGRGGSGGGGGGGGGGYNRSSGGYEPRGRGGGRG
GRGGMGGSDRGGFNKFGGPRDQGSRHDSEQDNSDNNTIFVQGLGENVTIESVADYFKQIGIIKTNKKTGQPMINLYTDRETGKL
KGEATVSFDDPPSAKAAIDWFDGKEFSGNPIKVSFATRRADFNRGGGNGRGGRGRGGPMGRGGYGGGGSGGGGRGGFPSGGG
GGGGQQRAGDWKCPNPTCENMNFSWRNECNQCKAPKPDGPGGGPGGSHMGGNYGDDRRGGRGGYDRGGYRGRGGDRGGF
RGGRGGGDRGGFGPGKMDSRGEHRQDRRERPY

The following Protein Data Bank (PDB) codes were used to build the globular structured domains of FUS (residues from 285–371 (PDB code: 2LCW) and from 422–453 (PDB code: 6G99)).

#### C. Colloidal patchy particle model for protein/RNA phase-separation

For the minimal coarse-grained simulations shown in Fig. 4 of the main text, we employ a patchy particle model [4–7] in which proteins are described by a pseudo hard-sphere (PHS) potential [8] that accounts for their excluded volume:

$$E_{PHS} = \sum_{i<j} \begin{cases} \lambda_r (\frac{\lambda_r}{\lambda_a})^{\lambda_a} \varepsilon_R \left[ (\frac{\sigma}{r_{ij}})^{\lambda_r} - (\frac{\sigma}{r_{ij}})^{\lambda_a} \right] + \varepsilon_R; & \text{if } r < (\frac{\lambda_r}{\lambda_a})\sigma \\ 0; & \text{if } r \geq (\frac{\lambda_r}{\lambda_a})\sigma \end{cases} . \tag{5}$$

where $\lambda_a$ =49 and $\lambda_r$=50 are the exponents of the attractive and repulsive terms respectively, and $\varepsilon_R$ accounts for the energy shift of the pseudo hard-sphere interaction. On top of this, we add a continuous square-well (CSW) potential for modeling the different protein binding sites, therefore mimicking protein multivalency:

$$E_{CSW} = \sum_{i<j} -\frac{1}{2}\epsilon_{CSW} \left[ 1 - tanh(\frac{r_{ij} - r_w}{\alpha}) \right] \tag{6}$$

where $\epsilon_{CSW}$ is the depth of the potential energy well, $r_w$ the radius of the attractive well, and $\alpha$ controls the steepness of the well. We choose $\alpha = 0.005\sigma$ and $r_w = 0.12\sigma$ so that each binding site can only interact with another single one. RNA-protein interactions are modeled with a standard Lennard-Jones (LJ) potential [9]:

$$E_{LJ} = \sum_{i<j} 4\epsilon_{LJ} \left[ \left( \frac{\sigma}{r_{ij}} \right)^{12} - \left( \frac{\sigma}{r_{ij}} \right)^{6} \right] \tag{7}$$

where $\epsilon_{LJ}$ measures the depth of potential and $\sigma$ the excluded volume between proteins and RNA. The Lennard-Jones potential is employed between protein cores (not binding sites) and RNA beads, while RNA-RNA interactions are modelled via the PHS potential [8], with a repulsive Yukawa potential on top of the PHS one:

$$E_{Yukawa} = \sum_{i<j} A \frac{e^{-\kappa r_{ij}}}{r_{ij}} \tag{8}$$

where we set A at 0.42 kcal·Å / mol and $\kappa$ at 2.57 Å$^{-1}$. In this way, we model RNA as a self-repulsive polymer of bonded repulsive spheres, and protein-RNA interactions via sites that are not the protein-protein binding sites. Hence, if one protein is bonded to RNA, it can still bind to other proteins. In our model, the mass of each patch is 5% of the central PHS particle mass, which is set to 3.32x10$^{-26}$kg, despite this choice being irrelevant for equilibrium simulations. This 5% ratio fixes the moment of inertia of the patchy particles (our minimal coarse-grained proteins). The molecular diameter of the proteins, both scaffold and cognate proteins, as well as the RNA beads is $\sigma = 0.3405$

nm, and the value of $\varepsilon_R/k_B$ is 119.81K. With this model, we express magnitudes in reduced units: reduced temperature is defined as $T^* = k_BT/\epsilon_{CSW}$, reduced density as $\rho^* = (N/V)\sigma^3$, reduced pressure as $p^* = p\sigma^3/(k_BT)$, and reduced time as $\sqrt{\sigma^2 m/(k_BT)}$. In order to keep the PHS interaction as similar as possible to a pure HS interaction, we fix $k_BT/\varepsilon_R$ at a value of 1.5 as suggested in Ref. [8] (fixing $T = 179.71$K). We then control the effective strength of the binding protein attraction by varying $\epsilon_{CSW}$ such that the reduced temperature, $T^* = k_BT/\epsilon_{CSW}$, is of the order of $\mathcal{O}(0.1)$. The cut-off distance for the interactions in this model are $1.17\sigma$ for both PHS and CSW potentials and $5\sigma$ for the LJ interactions. The $\epsilon_{LJ}/k_B$ for LJ interactions is set to 152.5K.

This model has been proven to qualitatively reproduce the effect of protein valency in LLPS [5], the enhancement of RNA-mediated LLPS in RNA-binding proteins [10] or the multilayer organization [6] and condensate size conservation in scaffold-client mixtures [7].

#### D. Simulation details

Our Direct Coexistence simulations [11, 12] are performed in the NVT ensemble (i.e. constant number of particles (N), volume (V) and temperature (T)), for which we use a Nosé–Hoover thermostat [13, 14] with a relaxation time of 5 ps for the Mpipi model simulations and 0.074 in reduced units for the patchy particle simulations. Since all our potentials are continuous and differentiable, we perform all our simulations using the LAMMPS Molecular Dynamics package[15]. Periodic boundary conditions are used in the three directions of space. The timestep chosen for the Verlet integration of the equations of motion is 10 fs for the Mpipi model and 3.7x$10^{-4}$ in reduced time units for the patchy particle model. For the bulk simulations with the Mpipi model, we employ the NpT ensemble, controlling the pressure (p) with the Nosé–Hoover barostat [16], with a relaxation time of 10 ps.

$PR_{25}$-polyU simulations were performed with 96 repeats of the protein and varying amounts of RNA, ranging from 1600 to 4800 nucleotides in total, and split into different chains to achieve the desired mass ratio and polyU strand length. For FUS-polyU simulations, we make use of 48 protein replicas and 400 to 1600 nucleotides in total forming RNA chains of different lengths. In the minimal model simulations we made use of 1000 coarse-grained proteins along with 100 to 600 RNA beads in total split into chains of different lengths as indicated in the main text.

### II. COMPUTING PHASE DIAGRAMS VIA DIRECT COEXISTENCE

To calculate the coexisting densities of the phase diagrams shown in Fig. 1b and 4b-c of the main text, we employ the Direct Coexistence method [11, 12, 17]. Within this scheme, the two coexisting phases are simulated by preparing periodically extended slabs of the two phases, the condensed and the diluted one, in the same simulation box. We use an implicit solvent model; accordingly, the diluted phase (protein-poor liquid phase) and the condensed phase (protein-rich liquid phase) are effectively a vapour and a liquid phase, respectively. Once our DC simulations have reached equilibrium, we compute the density profile along the long axis of the box, and thus, we extract the density of the two coexisting phases (as shown in Fig. 1(b) and 3(a-e) of the main text and the Supporting Material of Ref. [10]). From the plateau of the condensed phase and the diluted one, we measure the density (avoiding the interfaces between both phases). To estimate the critical point of the phase diagrams, we use the universal scaling law of coexistence densities near a critical point [18], and the law of rectilinear diameters [19]:

$$\big(\rho_l(T) - \rho_v(T)\big)^{3.06} = d\left(1 - \frac{T}{T_c}\right) \tag{9}$$

and

$$(\rho_l(T) + \rho_v(T))/2 = \rho_c + s_2(T_c - T) \tag{10}$$

where $\rho_l$ and $\rho_v$ refer to the coexisting densities of the condensed and diluted phases respectively, while $\rho_c$ is the critical density, $T_c$ is the critical temperature, and $d$ and $s_2$ are fitting parameters. On the other hand, the methodology to estimate critical temperatures via NpT bulk simulations is provided in Section II of the main text, and illustrated in Fig. 1b.

### III. COMPUTING THE INTERFACIAL FREE ENERGY FROM DIRECT COEXISTENCE SIMULATIONS

From Direct Coexistence simulations, we can obtain the interfacial free energy ($\gamma$) of the condensates by using the following expression:

$$\gamma = \int_{-\infty}^{\infty} [p_N(x) - p_T(x)]\, dx = \frac{L_x}{2N}(\overline{p}_N - \overline{p}_T) \tag{11}$$

where $L_x$ corresponds to the long side of the simulation box (perpendicular to the slab interfaces), $N$ is the number of droplets in the system, and $p_N$ and $p_T$ are the normal and tangential components of the pressure tensor with respect to the interfaces of the system (note that the tangential component must be averaged over the two tangential directions).


[1] A. J. Joseph, A. Reinhardt, A. Aguirre, P. Y. Chew, K. O. Russell, J. R. Espinosa, A. Garaizar, and R. Collepardo-Guevara, Nat. Comput. Sci. , in press (2021).
[2] P. Debye and E. Hückel, Physikalische Zeitschrift **24**, 185 (1923).
[3] X. Wang, S. Ramírez-Hinestrosa, J. Dobnikar, and D. Frenkel, Physical Chemistry Chemical Physics **22**, 10624 (2020).
[4] J. R. Espinosa, A. Garaizar, C. Vega, D. Frenkel, and R. Collepardo-Guevara, J. Chem. Phys **150**, 224510 (2019).
[5] J. R. Espinosa, J. A. Joseph, I. Sanchez-Burgos, A. Garaizar, D. Frenkel, and R. Collepardo-Guevara, Proceedings of the National Academy of Sciences (2020).
[6] I. Sanchez-Burgos, J. R. Espinosa, J. A. Joseph, and R. Collepardo-Guevara, Biomolecules **11**, 278 (2021).
[7] I. Sanchez-Burgos, J. A. Joseph, R. Collepardo-Guevara, and J. R. Espinosa, bioRxiv (2021).
[8] J. Jover, A. J. Haslam, A. Galindo, G. Jackson, and E. A. Müller, Journal of Chemical Physics **137** (2012), 10.1063/1.4754275.
[9] J. E. Jones, Proceedings of the Royal Society of London. Series A, Containing Papers of a Mathematical and Physical Character **106**, 441 (1924).
[10] J. A. Joseph, J. R. Espinosa, I. Sanchez-Burgos, A. Garaizar, D. Frenkel, and R. Collepardo-Guevara, Biophysical Journal **120**, 1219 (2021).
[11] A. J. Ladd and L. V. Woodcock, Chemical Physics Letters **51**, 155 (1977).
[12] R. García Fernández, J. L. F. Abascal, and C. Vega, The Journal of Chemical Physics **124**, 144506 (2006).
[13] S. Nosé, The Journal of Chemical Physics **81**, 511 (1984).
[14] W. G. Hoover, Phys. Rev. A **31**, 1695 (1985).
[15] S. Plimpton, Journal of Computational Physics **117**, 1 (1995).
[16] W. G. Hoover, Physical Review A **34**, 2499 (1986).
[17] J. R. Espinosa, E. Sanz, C. Valeriani, and C. Vega, Journal of Chemical Physics **139** (2013), 10.1063/1.4823499.
[18] J. S. Rowlinson and B. Widom, *Molecular theory of capillarity* (Courier Corporation, 2013).
[19] J. A. Zollweg and G. W. Mulholland, The Journal of Chemical Physics **57**, 1021 (1972).
